# Passive stomatal closure under extreme drought in an angiosperm species

**DOI:** 10.1101/2023.10.31.565010

**Authors:** Scott A. M. McAdam, Anju Manandhar, Cade N. Kane, Joel A. Mercado-Reyes

## Abstract

The phytohormone abscisic acid (ABA), synthesized as leaf turgor declines, plays a major role in closing stomata in species from this lineage, but recent reports of some angiosperms having a peaking-type ABA dynamic in which under extreme drought ABA levels decline to pre-stressed levels raises the possibility that passive stomatal closure by leaf water status alone can occur in species from this lineage. To test this hypothesis, we conducted instantaneous rehydration experiments in the peaking-type species *Umbellularia californica* through a long-term drought in which ABA levels declined to pre-stress levels yet stomata remain closed. We found that when ABA levels were lowest during extreme drought stomata of *U. californica* were passively closed by leaf water status alone, with stomata reopening rapidly to maximum rates of gas exchange on instantaneous rehydration. This contrasts with leaves early in drought in which ABA levels are highest, where we found stomata do not reopen on instantaneous rehydration. The transition from ABA driven stomatal closure to passively driven stomatal closure as drought progresses in this species occurs at very low water potentials facilitated by highly embolism resistant xylem. These results have important implications for understanding stomatal control during drought in angiosperms.

## Introduction

Stomata are mobile pores on the surface of leaves that regulate the exchange of water vapour from the plant with CO_2_ from the atmosphere (Raschke 1975). Unlike stomatal responses to light, CO_2_ and photosynthesis which optimize gas exchange such that photosynthesis is maximized for a given rate of water loss (Wong et al. 1979), the stomatal response to declining leaf water status restricts the majority of gas exchange with the atmosphere to conserve internal water reserves (Brodribb et al. 2017). This is because most plants are unable to survive tissue desiccation. The sensitivity of stomata to soil water content and the effectiveness of stomatal closure during drought are critical in determining the survival time of an individual when available soil water is depleted (Martin-StPaul et al. 2017). Considerable effort is placed on understanding the specific environmental or physiological drivers of stomatal closure when soil water content declines (Tardieu and Davies 1993; Verslues et al. 2022; Abdalla et al. 2021; Bourbia et al. 2021). In terms of how stomata respond to declining soil water status two mechanisms are recognized, the first is a direct, passive regulation of stomatal aperture by leaf water status (Rodriguez-Dominguez et al. 2016; Cardoso et al. 2019) and the other is an active hormonal trigger of stomatal closure, primarily through the hormone abscisic acid (ABA) (Tardieu and Simonneau 1998; McAdam and Brodribb 2012; Tombesi et al. 2015), which is synthesized when mesophyll cells begin to lose turgor (Pierce and Raschke 1980; Bacete et al. 2022).

The relative importance of passive or active hormonal driven processes on stomatal closure during drought vary across species, with the greatest difference being readily observed between seed and seed-free vascular land plants (McAdam and Brodribb 2014, 2012; Gong et al. 2021). In seed-free plants, stomatal closure during drought is passively driven by declines in leaf water status. Evidence for this simple, hydraulic regulation of stomatal closure during drought comes from observations of rapid stomatal reopening on instantaneous rehydration of leaves or whole plants when stomata are shut by drought (Cardoso et al. 2019; McAdam and Brodribb 2012). When rehydrated the stomata of species of seed-free plant reopen rapidly to maximum apertures as fast as the tissue rehydrates (Cardoso et al. 2019; McAdam and Brodribb 2012). This rapid reopening of stomata on the relaxation of leaf water potential provides compelling evidence for a passive mechanism driving stomatal closure because it occurs even when endogenous ABA levels in leaves are very high, indicating that endogenous levels of this hormone play no role in closing the stomata in species from these groups during drought (Cardoso and McAdam 2019). In contrast to seed-free plants, in species of seed plant in which the hormone ABA is synthesized during drought and levels are high in the leaf, similar rehydration experiments during drought fail to reopen stomata to maximum apertures (McAdam and Brodribb 2012; Brodribb and McAdam 2013). The degree of reopening of stomata in species of conifer, which do not have mechanical interactions between guard cells and epidermal cells, correlates with the level of ABA in the leaf at the time of rehydration (McAdam and Brodribb 2014). These experiments combined with observations in ABA biosynthetic and signalling mutants in angiosperm species which have stomata that remain largely open during drought (Xie et al. 2006; Tal and Nevo 1973), with mutant plants dying rapidly on soil water depletion (Brodribb et al. 2021), make a compelling case for the importance of hormonal regulation of plant water status in seed plants.

In some species of Cupressaceae and Taxaceae (Brodribb et al. 2014) and in extremely embolism resistant species of Fabaceae (Nolan et al. 2017; Yao et al. 2021b; Yao et al. 2021a) and Lauraceae (Mercado-Reyes et al. 2023) ABA levels increase as soil water potential begins to decline, closing stomata, but then once plants reach a water potential that approximates bulk leaf turgor loss point then ABA levels in the leaf begin to decline. Rapid rehydration experiments in the Cupressaceae conifer *Callitris* indicates that when ABA levels are low under extreme drought the stomata reopen to maximum apertures as fast as leaf water potential relaxes (Brodribb and McAdam 2013; McAdam and Brodribb 2015). These experiments demonstrate that in *Callitris* stomata transition from ABA driven stomatal closure early during a drought to a passive regulation of stomatal closure once bulk leaf turgor is lost and ABA levels decline. The recent description of a peaking-type ABA dynamic in some species of angiosperm suggests that passive stomatal closure during drought might also be found in species from all lineages of vascular land plants.

There are a few reasons why passive stomatal closure during drought in angiosperms is not widely accepted, the first is that mutants of ABA biosynthesis and signalling do not close effectively during drought or when leaf water status changes (Brodribb et al. 2021; Tulva et al. 2023), suggesting that if there is a passive regulation of stomatal response to leaf water status in species from this group of land plants it is minor or ineffective. Stomatal responses to short-term changes in leaf water status induced by vapour pressure difference between the leaf and the atmosphere (VPD) in angiosperms are not predictable by a passive-hydraulic model whereby guard cell turgor is linked to leaf turgor (Binstock et al. 2023; Cardoso et al. 2020). This contrasts with the stomatal responses to VPD in most non-angiosperm species (Brodribb and McAdam 2011). The close mechanical interactions between guard cells and epidermal cells which is unique to angiosperms and species of Marsileaceae (Westbrook and McAdam 2020), mean that stomatal aperture does not decline when leaf water status declines, at least not until epidermal turgor is lost (Buckley 2019, 2016). Finally, in species of *Caragana* (Fabaceae) in which a peaking type ABA dynamic during extreme drought has been reported, the gaseous hormone ethylene has been suggested to close stomata during drought once ABA levels have declined (Yao et al. 2021b).

To resolve whether the stomata of angiosperm species can be passively closed by leaf water status alone when soil water content declines, or whether species from this lineage always require a metabolic signal to promote stomatal closure we conducted instantaneous rehydration experiments on branches of *Umbellularia californica* (Lauraceae) during a long-term drought to investigate stomatal control when water deficit is removed. We have recently described this species as having a peaking-type ABA dynamic during drought, which means ABA levels are low and stomata are closed under extreme drought (Mercado-Reyes et al. 2023). We hypothesize that if stomata are closed passively under extreme drought in this angiosperm species, then on instantaneous rehydration stomata will rapidly reopen to maximum apertures.

## Materials and Methods

Five-year-old plants grown in 5 l pots containing a mix of Indiana Miami topsoil, ground pine bark and sand (1:2:1 ratio) were used for experiments. Plants were grown under controlled greenhouse conditions under a 16 h photoperiod (supplemented by LED lighting providing a PPFD of at least 150 μmol quanta m^−2^ s^−1^ at pot height), 28/22°C day/night temperature. Plants received daily irrigation when not under water deficit, and a monthly application of liquid fertilizer (Miracle-Gro® Water-Soluble All Purpose Plant Food, The Scotts Company LLC). Prior to imposing drought leaf gas exchange was measured using an infrared gas analyser (Li-6800, Licor, Lincoln NE) in leaves from three branches. Condition in the cuvette of the gas analyser were controlled to match the conditions in the greenhouse, with a chamber temperature set to 28°C, the leaf-to-air vapour pressure deficit set to 1.2 kPa, incoming air was drawn from a buffer drum so that CO_2_ in the cuvette matched ambient conditions in the greenhouse (approximately 420 µmol mol^-1^). Conditions in the cuvette were automatically logged every 30 s. Once acclimated to conditions in the cuvette and gas exchange was stable the branches were excised under water and gas exchange recorded until stable, this allowed us to recorded maximum stomatal conductance in hydrated branches excised under water, which allows us to account for any minor hydropassive stomatal closure that might occur on excision of angiosperm leaves under water (McAdam and Brodribb 2012). Once gas exchange was stable this value was used as the rate of stomatal conductance in unstressed branches that were fully hydrated. Leaf tissue was then harvested for both leaf water potential determination using a Scholander pressure chamber followed by the physicochemical quantification of ABA levels using an added internal standard according to the methods of Mercado-Reyes et al. (2023).

A soil water deficit was imposed on the plants by withholding water. Periodically during drought leaf water potential and foliage ABA levels were measured. Canopy conductance during the drought was determined gravimetrically by bagging pots and determining water loss rates over solar midday each day according to Mercado-Reyes et al. (2023). Once stomata had closed during drought, at a water potential at which ABA levels were known to be the highest during drought a leaf was again enclosed in the cuvette of the gas analyser (using the conditions described above). A neighbouring leaf was taken for the determination of leaf water potential and ABA levels. After leaf gas exchange had stabilized then the branch was excised under water and instantaneously rehydrated (occurring over less than 10 min). Leaf gas exchange was logged until gas exchange had again stabilized. The leaf in the cuvette was then taken for determining leaf water potential and foliage ABA levels. This experiment was repeated again once the levels of ABA in the leaf had declined to pre-stressed levels under long-term drought (approximately 25 d after stomatal closure).

## Results

In *U. californica* stomata close during a long-term drought (Figure 1A) and ABA levels display a classical peaking-type dynamic in which they increase early during drought then decline to prestressed levels as water potentials continue to decline (Figure 1A). When leaves are instantaneously rehydrated the degree of stomatal reopening was found to be associated with the level of ABA in the leaf (Figure 1). When ABA levels were high in a leaf, at 6.33 µg g^-1^ FW when the plant was at a water potential of -3.15 MPa and stomata were closed, instantaneous rehydration did not cause stomata to significantly open despite water potentials relaxing to -0.54 MPa in 15 min (Figure 1B). Once water potential had declined to -3.83 MPa, ABA levels had declined to 3.85 µg g^-1^ FW, an intermediate level between the peak of ABA levels and pre-stressed levels, and upon instantaneous rehydration, stomatal conductance rapidly increased to around 0.015 mol m^-2^ s^-1^ which was less than half of stomatal conductance in leaves from plants that had not been exposed to drought and in which ABA levels were low (Figure 1C). In branches in which ABA levels had declined to levels measured prior to drought (between 0.56-1.85 µg g^-1^ FW) instantaneous rehydration resulted in the rapid reopening of stomata to maximum conductance recorded in hydrated leaves measured in plants that had low levels of ABA prior to drought (Figures 1D-F). In all of these leaves leaf water potentials rapidly relaxed on rehydration (Figure 1D-F).

**Figure 1.**
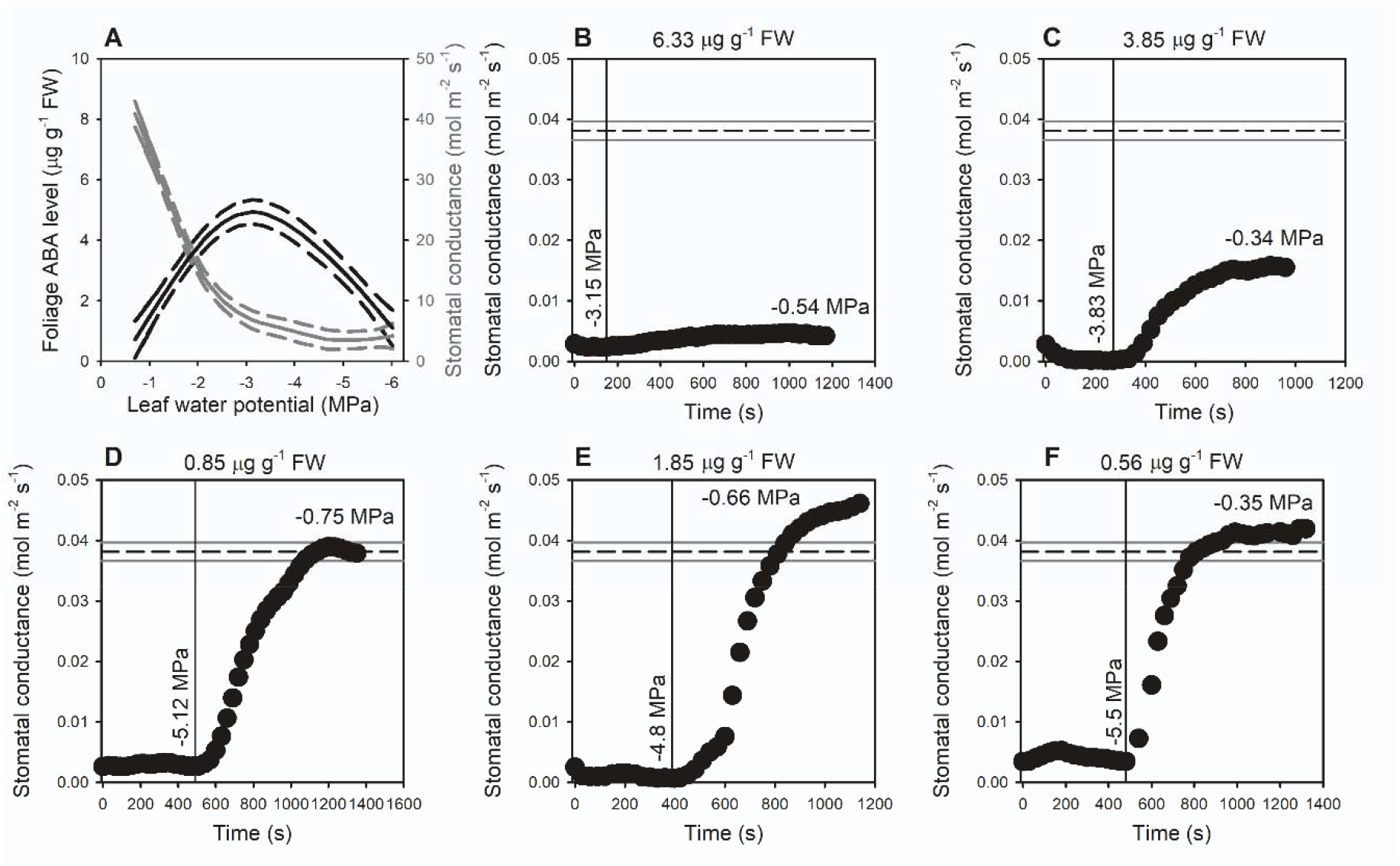
Stomata transition from ABA driven stomatal closure early during drought to passively driven stomatal closure under extreme drought in the angiosperm species *Umbellularia californica*. (A) The relationship between stomatal conductance (grey) and foliage ABA level (black) during drought in *U. californica*, data are generalized additive models (±SE) taken from Mercado-Reyes et al. (2023) from the same plants from which gas exchange data was collected. (B-F) The response of stomatal conductance (black dots) to instantaneous branch rehydration (denoted by the vertical line), the foliage ABA level is shown above each panel, the water potential prior to rehydration is shown parallel to the line denoting rehydration, the water potential at the end of the gas exchange trace is also shown. The mean (n=5, ±SE) maximum stomatal conductance in unstressed branches that are excised under water with a mean ABA level of 0.93 ±0.02 µg g-1 FW is shown as a vertical dashed line bound by the standard error.

## Discussion

We show here that the stomata of the extremely drought tolerant angiosperm species *U. californica* which has a leaf P_50_ (or water potential at which 50% of xylem is embolized) of -7.54 MPa (Mercado-Reyes et al. 2023), transitions to passive stomatal closure, from closure driven by the hormone ABA under long term drought (Figure 1). Our simple experiment in which stomatal conductance was recorded during instantaneous rehydration of branches taken from drought stressed plants demonstrates that high levels of ABA in the leaf early in a drought are sufficient to keep stomata closed on rapid rehydration. This is similar to observations made across seed plant species which synthesize high levels of ABA during drought and that high levels of endogenous ABA keeps stomata closed in seed plants (McAdam and Brodribb 2012, 2014). We found that once ABA levels have declined to pre-stressed levels when plants approached -5 MPa, stomata become passively closed by low leaf water status alone. That the stomata of *U. calfornica* under these conditions could reopen rapidly to a conductance recorded in unstressed leaves suggests that neither ABA nor any other hormone or metabolic signal is keeping stomata closed during extreme drought in this species. This challenges the view that ethylene might be responsible for driving stomatal closure during drought in angiosperm species in which a peaking-type ABA dynamic has been observed (Yao et al. 2021b).

Our observations in *U. californica* are very similar to work in the genus *Callitris* in which instantaneous rehydration during a drought when ABA levels are lowest results in stomata reopening to maximum conductance (Brodribb and McAdam 2013). It has been hypothesized that this ability to rapidly recover maximum rates of gas exchange is advantageous for species that grow in environments that grow in seasonally dry environments, allowing for the rapid and uninhibited recovery of photosynthetic capacity following rainfall (McAdam and Brodribb 2015). We also observed that as ABA levels started to decline during drought that the residual ABA continues to keep stomata closed, with stomata not reopening to maximum conductance on instantaneous rehydration when ABA levels were half the level measured in leaves when at the peak early in drought. This also mirrors work in *Callitris* in which there is a transition period between stomatal closure being exclusively driven by ABA to being driven by both low water status and ABA levels, to finally just by low leaf water status alone (Brodribb and McAdam 2013). Our results support a theory that ABA promotes early stomatal closure during drought but that as leaf water status declines sufficiently then stomata can be closed passively in angiosperms (Rodriguez-Dominguez et al. 2016). This differs from work in grape which suggests that declining water status passively closes stomata early in a drought and that ABA keeps stomata closed under long-term drought, although this conclusion comes from work on a species with much more vulnerable xylem to embolism formation (Tombesi et al. 2015). Our work also suggests that the capacity to sustain high turgor pressures in guard cells in the light is balanced by low plant water status. By showing that stomata could reopen rapidly to maximum conductance in the light when instantaneously rehydrated after more than 25 days of being closed during drought indicates that the capacity to actively load ions into the guard cells in the light (via the H+-ATPase) is not inhibited by low leaf water potential when ABA is not present (Pei et al. 2022).

All data collected to date that reports a peaking-type ABA dynamic during drought has found that this phenomenon only occurs at very low water potentials (less than -4 MPa) and consequently a requirement is highly embolism resistant xylem (Brodribb et al. 2014; Nolan et al. 2017; Yao et al. 2021b; Yao et al. 2021a; Mercado-Reyes et al. 2023). Recent work in *U. californica* and *C. rhomboidea* suggests that ABA biosynthesis is deactivated at water potentials lower than turgor loss point and requires sufficient time for the residual ABA to be conjugated for the levels to decline (Mercado-Reyes et al. 2023). Modelling suggests that passive stomatal control in angiosperms is possible once epidermal turgor is lost and the mechanical advantage of the epidermis is removed (Buckley 2019). If epidermal turgor loss is assumed to occur at the same leaf water potential as the loss of bulk leaf turgor, then for most herbaceous plants in which embolism in the xylem forms at water potentials close to turgor loss point, leaf death would supersede any observations of declines in ABA levels associated with passively closed stomata (Skelton et al. 2017). A key question that remains unanswered but emerges from our dataset is whether the stomata of angiosperms can be passively closed by declines in leaf water status when soil water status is high, for example when VPD increases (Merilo et al. 2017)? There are reports from single gene mutants in key ABA signalling and synthesis genes that suggest there is a small degree of stomatal response to VPD that might be due to a passive response in herbaceous angiosperms (Merilo et al. 2017), however there are also conflicting reports from multi-order mutants that suggest these responses in single-gene mutants might be due to allelic leakiness or genetic redundancy in the pathways of ABA synthesis and response, and that when ABA signalling and synthesis is completely blocked stomata do not respond to VPD (Brodribb et al. 2021; Fujii et al. 2011). We do not yet know what maximum turgor pressures are in the guard cells of open stomata in angiosperms compared to the stomata of other lineages of land plants and whether passive declines in leaf water status driven by high VPD would be sufficient to appreciably change the aperture of the pore, like they are hypothesized to do in species of fern and lycophyte (Franks and Farquhar 2007; Brodribb and McAdam 2011).

Here we show that stomata of a highly embolism resistant angiosperm species can be closed passively by leaf water status alone under an extreme drought. Our results suggest that early during drought stomata are closed by ABA but that they transition to being closed by leaf water status alone when ABA levels being to decline. Our observations of rapid stomatal reopening on instantaneous rehydration from a state of stomatal closure at very low water potentials suggests that the stomata of species from across all lineages of vascular land plant have the potential to be passively regulated. Whether this passive regulation occurs at mild water potentials more akin to those experienced by herbaceous plants under drought, or when VPD increases in angiosperms remains to be tested.

## Acknowledgements

We thank Missy Holbrook for helpful discussions. We acknowledge the use of the Metabolite Profiling Facility of the Bindley Bioscience Center, a core facility of the NIH-funded Indiana Clinical and Translational Sciences Institute for quantifying hormone levels.

## Author contributions

SM designed the experiments and wrote the manuscript with the help of AM and CK; SM, CK and JMR collected the data.

## Data availability

All data are available on request from the authors.

## References

Abdalla M, Ahmed MA, Cai G, Wankmüller F, Schwartz N, Litig O, Javaux M, Carminati A (2021) Stomatal closure during water deficit is controlled by below-ground hydraulics. Annals of Botany 129 (2):161–170. doi:10.1093/aob/mcab141

Bacete L, Schulz J, Engelsdorf T, Bartosova Z, Vaahtera L, Yan G, Gerhold JM, Tichá T, Øvstebø C, Gigli-Bisceglia N, Johannessen-Starheim S, Margueritat J, Kollist H, Dehoux T, McAdam SAM, Hamann T (2022) THESEUS1 modulates cell wall stiffness and abscisic acid production in Arabidopsis thaliana. Proceedings of the National Academy of Sciences 119 (1):e2119258119. doi:doi:10.1073/pnas.2119258119

Binstock BR, Manandhar A, McAdam SAM (2023) Characterizing the breakpoint of stomatal response to vapor pressure deficit in an angiosperm. Plant Physiol. doi:10.1093/plphys/kiad560

Bourbia I, Pritzkow C, Brodribb TJ (2021) Herb and conifer roots show similar high sensitivity to water deficit. Plant Physiology 186 (4):1908–1918. doi:10.1093/plphys/kiab207

Brodribb T, Brodersen CR, Carriqui M, Tonet V, Rodriguez Dominguez C, McAdam S (2021) Linking xylem network failure with leaf tissue death. New Phytologist 232 (1):68–79. doi:10.1111/nph.17577

Brodribb TJ, McAdam SA, Carins Murphy MR (2017) Xylem and stomata, coordinated through time and space. Plant, Cell & Environment 40 (6):872–880. doi:10.1111/pce.12817

Brodribb TJ, McAdam SAM (2011) Passive origins of stomatal control in vascular plants. Science 331 (6017):582–585

Brodribb TJ, McAdam SAM (2013) Abscisic acid mediates a divergence in the drought response of two conifers. Plant Physiology 162:1370–1377

Brodribb TJ, McAdam SAM, Jordan GJ, Martins SCV (2014) Conifer species adapt to low-rainfall climates by following one of two divergent pathways. Proceedings of the National Academy of Sciences of the United States of America 111 (40):14489–14493. doi:10.1073/pnas.1407930111

Buckley TN (2016) Stomatal responses to humidity: has the “black box” finally been opened? Plant Cell Environ 39:482–484. doi:10.1111/pce.12651

Buckley TN (2019) How do stomata respond to water status? New Phytologist 224 (1):21–36. doi:10.1111/nph.15899

Cardoso AA, Brodribb TJ, Kane CN, DaMatta FM, McAdam SAM (2020) Osmotic adjustment and hormonal regulation of stomatal responses to vapour pressure deficit in sunflower. AoB PLANTS 12 (4). doi:10.1093/aobpla/plaa025

Cardoso AA, McAdam SAM (2019) Misleading conclusions from exogenous ABA application: a cautionary tale about the evolution of stomatal responses to changes in leaf water status. Plant Signaling & Behavior 14 (7):1610307. doi:10.1080/15592324.2019.1610307

Cardoso AA, Randall JM, McAdam SAM (2019) Hydraulics regulate stomatal responses to changes in leaf water status in the fern Athyrium filix-femina. Plant Physiology 179 (2):533–543. doi:10.1104/pp.18.01412

Franks PJ, Farquhar GD (2007) The mechanical diversity of stomata and its significance in gas-exchange control. Plant Physiology 143 (1):78–87

Fujii H, Verslues PE, Zhu JK (2011) Arabidopsis decuple mutant reveals the importance of SnRK2 kinases in osmotic stress responses in vivo. Proc Natl Acad Sci U S A 108 (4):1717–1722. doi:10.1073/pnas.1018367108

Gong L, Liu X-D, Zeng Y-Y, Tian X-Q, Li Y-L, Turner NC, Fang X-W (2021) Stomatal morphology and physiology explain varied sensitivity to abscisic acid across vascular plant lineages. Plant Physiology 186 (1):782–797. doi:10.1093/plphys/kiab090

Martin-StPaul N, Delzon S, Cochard H (2017) Plant resistance to drought depends on timely stomatal closure. Ecology Letters 20 (11):1437–1447. doi:10.1111/ele.12851

McAdam SAM, Brodribb TJ (2012) Fern and lycophyte guard cells do not respond to endogenous abscisic acid. The Plant Cell 24:1510–1521

McAdam SAM, Brodribb TJ (2014) Separating active and passive influences on stomatal control of transpiration. Plant Physiology 164:1578–1586

McAdam SAM, Brodribb TJ (2015) Hormonal dynamics contributes to divergence in seasonal stomatal behaviour in a monsoonal plant community. Plant, Cell and Environment 38 (3):423–432. doi:10.1111/pce.12398

Mercado-Reyes J, Pereira TS, Manandhar A, Rimer IM, McAdam SAM (2023) Extreme drought can deactivate ABA biosynthesis in embolism resistant species. Plant, Cell & Environment doi: 10.1111/pce.14754

Merilo E, Yarmolinsky D, Jalakas P, Parik H, Tulva I, Rasulov B, Kilk K, Kollist H (2017) Stomatal VPD Response: There Is More to the Story Than ABA Plant Physiology 176 (1):851–864. doi:10.1104/pp.17.00912

Nolan RH, Tarin T, Santini NS, McAdam SAM, Ruman R, Eamus D (2017) Differences in osmotic adjustment, foliar abscisic acid dynamics, and stomatal regulation between an isohydric and anisohydric woody angiosperm during drought. Plant Cell Environ 40 (12):3122–3134. doi:10.1111/pce.13077

Pei D, Hua D, Deng J, Wang Z, Song C, Wang Y, Wang Y, Qi J, Kollist H, Yang S, Guo Y, Gong Z (2022) Phosphorylation of the plasma membrane H+-ATPase AHA2 by BAK1 is required for ABA-induced stomatal closure in Arabidopsis. The Plant Cell 34 (7):2708–2729. doi:10.1093/plcell/koac106

Pierce M, Raschke K (1980) Correlation between loss of turgor and accumulation of abscisic acid in detached leaves. Planta 148 (2):174–182. doi:10.1007/BF00386419

Raschke K (1975) Stomatal action. Annu Rev Plant Physiol 26:309–340

Rodriguez-Dominguez CM, Buckley TN, Egea G, de Cires A, Hernandez-Santana V, Martorell S, Diaz-Espejo A (2016) Most stomatal closure in woody species under moderate drought can be explained by stomatal responses to leaf turgor. Plant, Cell & Environment 39 (9):2014–2026. doi:10.1111/pce.12774

Skelton RP, Brodribb TJ, Choat B (2017) Casting light on xylem vulnerability in an herbaceous species reveals a lack of segmentation. New Phytologist 214 (2):561–569. doi:10.1111/nph.14450

Tal M, Nevo Y (1973) Abnormal stomatal behavior and root resistance, and hormonal imbalance in three wilty mutants of tomato. Biochemical Genetics 8 (3):291–300. doi:10.1007/bf00486182

Tardieu F, Davies WJ (1993) Integration of hydraulic and chemical signalling in the control of stomatal conductance and water status of droughted plants. Plant Cell Environ 16:341–349

Tardieu F, Simonneau T (1998) Variability among species of stomatal control under fluctuating soil water status and evaporative demand: modelling isohydric and anisohydric behaviours. Journal of Experimental Botany 49 (1):419–432. doi:10.1093/jexbot/49.suppl_1.419

Tombesi S, Nardini A, Frioni T, Soccolini M, Zadra C, Farinelli D, Poni S, Palliotti A (2015) Stomatal closure is induced by hydraulic signals and maintained by ABA in drought-stressed grapevine. Sci Rep 5:12449. doi:10.1038/srep12449

Tulva I, Välbe M, Merilo E (2023) Plants lacking OST1 show conditional stomatal closure and wildtype-like growth sensitivity at high VPD. Physiologia Plantarum 175 (5):e14030. doi:10.1111/ppl.14030

Verslues PE, Bailey-Serres J, Brodersen C, Buckley TN, Conti L, Christmann A, Dinneny JR, Grill E, Hayes S, Heckman RW, Hsu P-K, Juenger TE, Mas P, Munnik T, Nelissen H, Sack L, Schroeder JI, Testerink C, Tyerman SD, Umezawa T, Wigge PA (2022) Burning questions for a warming and changing world: 15 unknowns in plant abiotic stress. The Plant Cell 35 (1):67–108. doi:10.1093/plcell/koac263

Westbrook AS, McAdam SAM (2020) Stomatal density and mechanics are critical for high productivity: insights from amphibious ferns. New Phytologist 229:877–889. doi:10.1111/nph.16850

Wong SC, Cowan IR, Farquhar GD (1979) Stomatal conductance correlates with photosynthetic capacity. Nature 282 (5737):424–426

Xie X, Wang Y, Williamson L, Holroyd GH, Tagliavia C, Murchie E, Theobald J, Knight MR, Davies WJ, Leyser HMO, Hetherington AM (2006) The identification of genes involved in the stomatal response to reduced atmospheric relative humidity. Current Biology 16 (9):882–887. doi:10.1016/j.cub.2006.03.028

Yao G-Q, Nie Z-F, Turner NC, Li F-M, Gao T-P, Fang X-W, Scoffoni C (2021a) Combined high leaf hydraulic safety and efficiency provides drought tolerance in *Caragana* species adapted to low mean annual precipitation. New Phytologist 229 (1):230–244. doi:10.1111/nph.16845

Yao GQ, Li FP, Nie ZF, Bi MH, Jiang H, Liu XD, Wei Y, Fang XW (2021b) Ethylene, not ABA, is closely linked to the recovery of gas exchange after drought in four *Caragana* species. Plant Cell Environ 44 (2):399–411. doi:10.1111/pce.13934

